# Comparative analysis of transcriptomic liberality in 2,3,7,8-tetrachlorodibenzo-*p*-dioxin–exposed mouse liver RNA-Seq datasets

**DOI:** 10.64898/2026.07.30.741944

**Authors:** Yuko Ogata, Kenichi Kobayashi

## Abstract

Omics methods have been envisioned to complement traditional toxicological testing for chemical risk assessment, in which identifying adverse effects is a critical step. However, the high dimensionality of transcriptomic data has historically led to reliance on context-dependent analysis. Liberality is a quantitative index that reduces genome-scale data dimensionality, with its changes reflecting underlying biological phenomena. In this study, we measured liberality in mouse liver RNA sequencing (RNA-Seq) datasets from studies in which mice were exposed to the environmental contaminant 2,3,7,8-tetrachlorodibenzo-*p*-dioxin (TCDD) every 4 days for 28 or 92 days, comparing dose–liberality relationships. Both 28- and 92-day TCDD treatments increased liberality but exhibited different dose–liberality relationships. Analysis of genes contributing to liberality revealed that longer exposure duration induced more extensive alterations in transcriptomic architecture. These findings suggest that liberality may serve as an unbiased metric to assess the extent of treatment-induced transcriptome perturbation.

## Introduction

Characterizing a treatment-induced effect as adverse or nonadverse is crucial in chemical risk assessment. Historically, this has been based on comparing clinical and/or pathological observations between chemically treated and control animals. Omics methods, part of New Approach Methodologies, have long been envisioned to complement traditional toxicological testing relying mainly on high-dose phenotypic response assessment in animals, because their comprehensive nature enables the investigation of molecular events underlying toxic responses and their sequential progression (Frohlich, 2017; Heijne *et al*., 2005; Vinken, 2019). However, this advantage is accompanied by the high dimensionality of Omics data, which makes analysis and interpretation more challenging than those of conventional toxicological test (Gant *et al*., 2023). To address this, previous studies have focused on identifying molecular features linked to apical outcomes or known modes of action within transcriptome datasets comprising hundreds or thousands of endpoints. Consequently, gene subsets relevant to toxicology (Li *et al*., 2015; Pan *et al*., 2025; Podtelezhnikov *et al*., 2020) or exhibiting clear exposure–response relationships under benchmark dose (Farmahin *et al*., 2017; Kawamoto *et al*., 2017; Thomas *et al*., 2007; Thomas *et al*., 2013) or benchmark concentration (Gao *et al*., 2024; Ramaiahgari *et al*., 2019) modeling frameworks have been selected and evaluated. In contrast, Barutcu *et al*. (2023) proposed a comprehensive pathway-based benchmark dose analysis without *a priori* selection to provide a complementary system-level perspective on chemical-induced biological perturbations.

Liberality, a quantitative measure derived from RNA sequencing (RNA-Seq) data, provides a means of reducing the dimensionality of genome-scale data and quantifies intra-sample diversity (Ogata, 2024; Ogata *et al*., 2012). Following the initial demonstration of its relationship with cellular differentiation and dedifferentiation (Ogata *et al*., 2012), liberality has been linked to a wide range of biological phenomena (Antolovic *et al*., 2019; Brehme *et al*., 2016; Ogata, 2024, 2025b; Ogata and Hosaka, 2022; Ogata *et al*., 2015; Tu *et al*., 2026; Wiesner *et al*., 2018). A previous study using human RNA-Seq data showed that diseased samples consistently exhibited higher liberality than healthy samples (Ogata, 2025a). Similar increases have been observed in data from mammalian models of liver disease (unpublished data). Based on these observations, liberality is expected to increase in the transcriptomes of organs adversely affected by chemical exposure.

This study aimed to investigate how liberality changes with exposure dose and duration, two factors closely associated with the severity of treatment-induced effects. Two previously published mouse liver RNA-Seq datasets from studies where mice were exposed to the environmental contaminant 2,3,7,8-tetrachlorodibenzo- *p*-dioxin (TCDD) every 4 days for 28 days (Nault *et al*., 2015) or 92 days (Nault *et al*., 2016) were selected, and their dose–liberality relationships were compared. We also discuss the potential utility of liberality for supporting the integration of transcriptomics into chemical risk assessment.

## Materials and Methods

### Data collection and analysis

We used publicly available count data from the National Centre for Biotechnology Information (NCBI) Gene Expression Omnibus (GEO) database. Liberality is described using an adaptation of Shannon’s entropy formula:

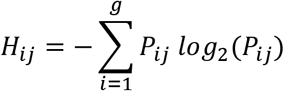

The relative expression level of gene *i* in tissue *j* is denoted by *P*_*ij*_. The following R code from previous studies (Ogata, 2024; Ogata *et al*., 2015) was used for its calculation:

entroshannon<-function(x){

x2 <- x/sum(x)

x3 <- x2*log2(x2)

x4 <- x3[!is.na(x3)]

-sum(x4)

}

To evaluate the dose–liberality relationship, linear (b ∼ a) and quadratic (b ∼ a + I(a^2)) regression models were fitted using the following R code, where “ie” represents the liberality value calculated using the entroshannon function described above:

a <- log(data$dose+0.00001)

b <- data$ie

c <- cbind(a, b)

model <- lm(b ∼ a)

model2 <- lm(b ∼ a + I(a^2))

The Akaike information criterion (AIC) was calculated for each model using R to compare model fit. Comparisons between linear and quadratic regression models were performed using the anova() function in R.

### Dataset description: 28-day repeated TCDD exposure

Nault *et al*. (2015) sequenced female C57BL/6 mouse liver transcriptomes. The mice received 0.1 mL sesame oil as a vehicle control or TCDD at doses of 0.01, 0.03, 0.1, 0.3, 1, 3, 10, or 30 µg/kg by oral gavage every 4 days for 28 days. Each dose group included five biological replicates.

We used RNA-Seq count data deposited under the following accession numbers: GSM1535917, GSM1535918, GSM1535919, GSM1535920, GSM1535921, GSM1535922, GSM1535923, GSM1535924, GSM1535925, GSM1535926, GSM1535927, GSM1535928, GSM1535929, GSM1535930, GSM1535931, GSM1535932, GSM1535933, GSM1535934, GSM1535935, GSM1535936, GSM1535937, GSM1535938, GSM1535939, GSM1535940, GSM1535941, GSM1535942, GSM1535943, GSM1535944, GSM1535945, GSM1535946, GSM1535947, GSM1535948, GSM1535949, GSM1535950, GSM1535951, GSM1535952, GSM1535953, GSM1535954, GSM1535955, GSM1535956, GSM1535957, GSM1535958, GSM1535959, GSM1535960, and GSM1535961. Prior to analysis, rows corresponding to no_feature, ambiguous, too_low_aQual, not_aligned, and alignment_not_unique in the generated count table were removed.

### Dataset description: 92-day repeated TCDD exposure

Nault *et al*. (2016) sequenced female C57BL/6 mouse liver transcriptomes. The mice received 0.1 mL sesame oil as a vehicle control or TCDD at doses of 0.01, 0.03, 0.1, 0.3, 1, 3, 10, or 30 µg/kg by oral gavage every 4 days for 92 days. Each dose group included three biological replicates.

We used RNA-Seq count data deposited under the following accession numbers: GSM2179711, GSM2179712, GSM2179713, GSM2179714, GSM2179715, GSM2179716, GSM2179717, GSM2179718, GSM2179719, GSM2179720, GSM2179721, GSM2179722, GSM2179723, GSM2179724, GSM2179725, GSM2179726, GSM2179727, GSM2179728, GSM2179729, GSM2179730, GSM2179731, GSM2179732, GSM2179733, GSM2179734, GSM2179735, GSM2179736, and GSM2179737. Prior to analysis, rows corresponding to no_feature, ambiguous, too_low_aQual, not_aligned, and alignment_not_unique in the generated count table were removed.

### Use of artificial intelligence tools

ChatGPT (v5.5; OpenAI, San Francisco, CA, USA) was used to help identify NCBI GEO accession numbers for relevant datasets and to locate some references cited in the present study. The authors verified all information obtained through ChatGPT against the original data sources and primary literature.

## Results and Discussion

Figure 1 illustrates the liberality–TCDD dose relationship for the 28-day (Fig. 1A) or 92-day (Fig. 1B) treatment. For the 28-day treatment, the measured liberalities were 10.07856, 10.1183, 10.14874, 10.08875, and 10.1813 for 0 µg/kg TCDD; 10.08685, 10.24628, 9.962552, 10.0137, and 10.22344 for 0.01 µg/kg TCDD; 10.1009, 10.21077, 10.17972, 10.09973, and 10.15161 for 0.03 µg/kg TCDD; 10.10825, 10.25743, 10.28774, 10.36162, and 10.03738 for 0.1 µg/kg TCDD; 10.27539, 10.19695, 10.24482, 10.29055, and 10.17179 for 0.3 µg/kg TCDD; 10.24434, 10.23118, 10.27938, 10.32904, and 10.13587 for 1 µg/kg TCDD; 10.27062, 10.34093, 10.28149, 10.29912, and 10.17323 for 3 µg/kg TCDD; 10.16314, 10.15125, 10.2914, 10.20051, and 10.16153 for 10 µg/kg TCDD; and 10.35118, 10.36677, 10.25499, 10.27858, and 9.884903 for 30 µg/kg TCDD. The slope of the linear regression model was 0.009417 and significantly differed from zero (p = 0.0103). The linear model’s AIC was -75.86214. The quadratic model’s AIC was −73.89914. Likelihood ratio testing showed no significant improvement in model fit with the quadratic term (p = 0.8534), indicating that the linear model adequately described the relationship.

**Figure 1.**
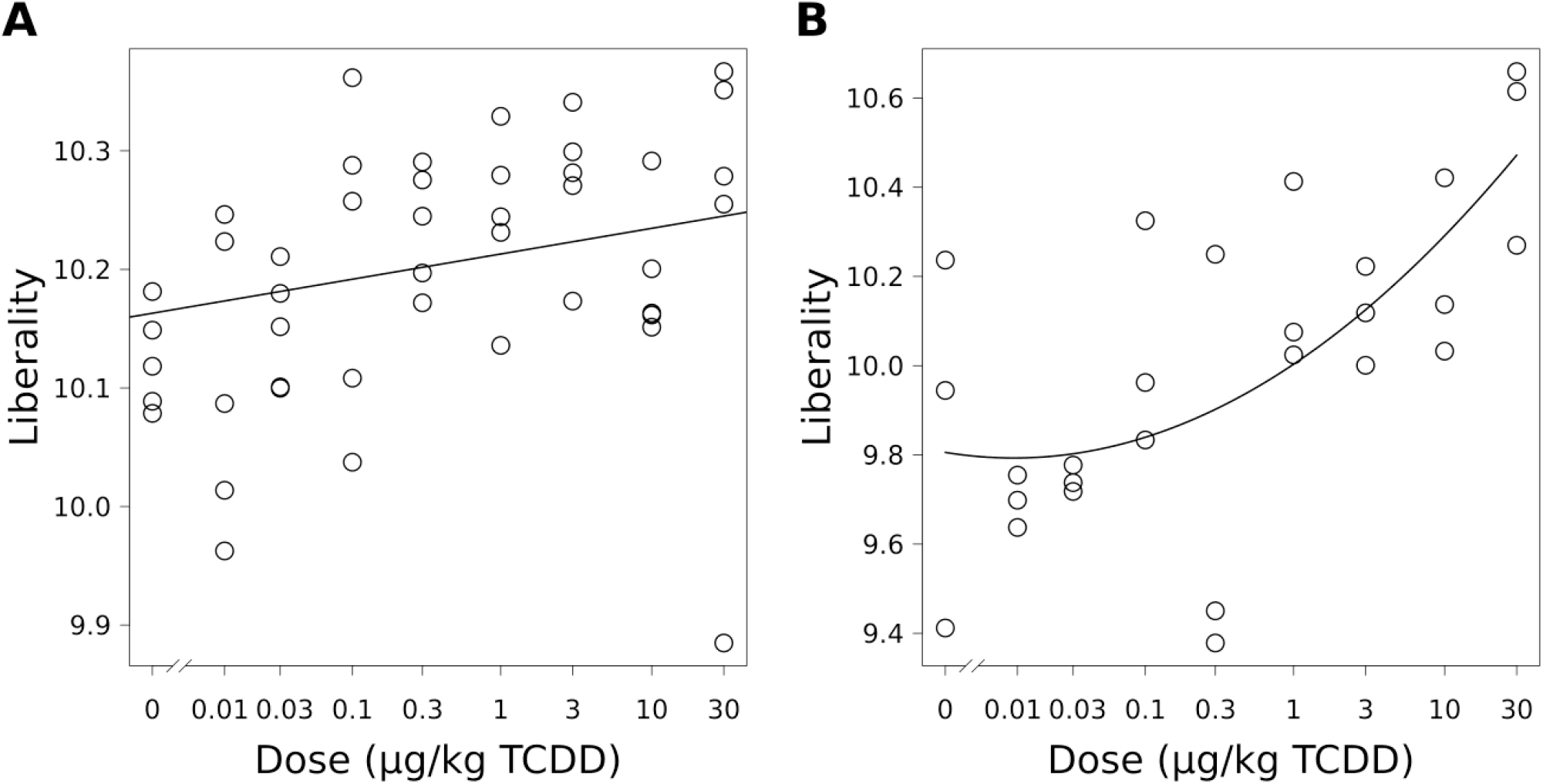
Scatter plots of TCDD dose versus liberality. (A) 28-day treatment. (B) 92-day treatment. TCDD: 2,3,7,8-tetrachlorodibenzo-*p*-dioxin.

Regarding the 92-day treatment, the observed liberalities were 10.23662, 9.944463, and 9.411965 for 0 µg/kg TCDD; 9.637348, 9.698411, and 9.754641 for 0.01 µg/kg TCDD; 9.737794, 9.71804, and 9.777132 for 0.03 µg/kg TCDD; 9.962249, 10.32489, and 9.833437 for 0.1 µg/kg TCDD; 10.24963, 9.378304, and 9.4504 for 0.3 µg/kg TCDD; 10.41288, 10.02416, and 10.07526 for 1 µg/kg TCDD; 10.00082, 10.11838, and 10.22247 for 3 µg/kg TCDD; 10.42109, 10.03263, and 10.13697 for 10 µg/kg TCDD; and 10.61465, 10.26984, and 10.65918 for 30 µg/kg TCDD. The linear regression slope was 0.04045 and significantly differed from zero (p = 0.0071). The AIC was 16.142042 for the linear model and 8.788426 for the quadratic model. Likelihood ratio testing indicated that the quadratic model better explained the dose–liberality relationship for 92 days compared with the linear model (*p* = 0.004311).

These results indicate that both treatments increased liberality dose-dependently. High entropy indicates a broader transcriptome diversity. Notably, liberality increased linearly with dose for 28 days but nonlinearly for 92 days. In the 92-day study, treatment-related losses occurred after 28 days; specifically, animals that exhibited a sudden ∼15% body weight decrease were subsequently terminated, whereas no overt toxicity was observed in the 28-day study (Nault *et al*., 2016). The findings in the 92-day study may indicate the systemic adverse effects of TCDD on the general health of mice. Nault *et al*. (2016) also reported the emergence of histopathological features of hepatic nonalcoholic fatty liver disease at lower doses in the 92-day study than in the 28-day study. Furthermore, an additional histopathological feature, extramedullary hematopoiesis, was observed in survivors treated with 30 µg/kg TCDD for 92 days. Unlike hepatic lipid accumulation, inflammation, and fibrosis, extramedullary hematopoiesis is generally regarded as a compensatory response to hematopoietic stress arising from conditions such as toxicant exposure, infection and inflammation (Cenariu *et al*., 2021; Johns and Christopher, 2012; Kim, 2010; NTP).

To explore the factors underlying the increase in liberality, we analyzed log_2_(P_ij_), a term in its formula, for genes contributing to liberality in the 10 and 30 µg/kg TCDD treatment groups, which exhibited high liberality values. The distribution of log_2_(P_ij_) for individual genes is shown in Fig. 2. This analysis revealed two key findings. First, TCDD exposure increased the number of genes contributing to liberality. The mean total number of genes with log_2_(P_ij_) values between −16.0 and −12.0 increased with dose: 5598, 5912.4, and 6089.6 for 28 days, and 5755.667, 6304.333, and 6685.333 for 92 days at 0, 10, and 30 µg/kg TCDD, respectively. Second, as a notable feature of 30 µg/kg TCDD exposure for 92 days, the log_2_(P_ij_) values of top-ranked genes (located at the top right in Fig. 2D) became similar. In this group, the relative expression level of albumin (ENSMUSG00000029368), which was consistently ranked first across samples, decreased, whereas expression levels of genes such as *Cyp1a2* (ENSMUSG00000032310) and NADH–ubiquinone oxidoreductase chain 1 (ENSMUSG00000064341) increased. The contribution of the 30 µg/kg treatment group to the nonlinearity of the dose–liberality relationship observed in the 92-day study is supported by AIC and likelihood ratio tests using the models described above. Specifically, when analyzing the 92-day dataset without the 30 µg/kg treatment group, AIC values were 12.424316 and 9.727493 for the linear and quadratic models, respectively (p = 0.04511). The increase in the number of expressed genes is consistent with previous studies indicating that the progression of TCDD-induced liver injury in mice is accompanied by the progressive involvement of multiple biological pathways (Cholico *et al*., 2024; Fader *et al*., 2015; Fling *et al*., 2020; Nault *et al*., 2016; Nault *et al*., 2023). The flattening of the expression levels of highly expressed genes observed in the 30 µg/kg TCDD group for 92 days may reflect a shift in cellular resource allocation in response to multiple concurrent demands.

**Figure 2.**
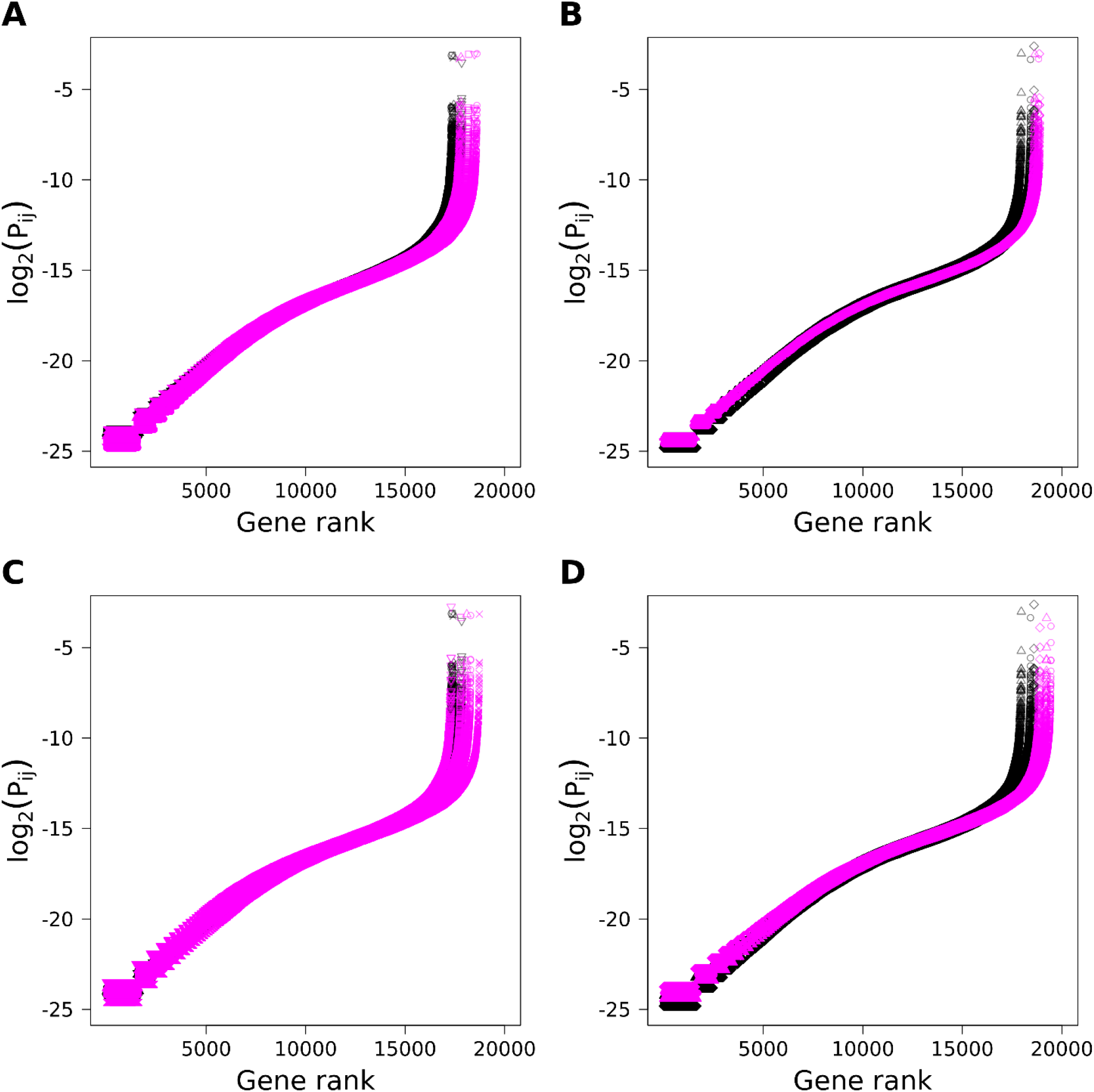
Log_2_(P_ij_) values of genes contributing to liberality in control and TCDD-treated groups. Genes contributing to liberality (x-axis) were ordered by read count, with low- and high-count genes on the left and right, respectively. Control samples: black; treated samples: pink. Genes with zero read counts in each sample were excluded. (A) Control vs. 10 µg/kg TCDD for 28 days. (B) Control vs. 10 µg/kg TCDD for 92 days. (C) Control vs. 30 µg/kg TCDD for 28 days. (D) Control vs. 30 µg/kg TCDD for 92 days.

Previous correlation analyses of fold-change and posterior probability P1(t) values for the 2,233 differentially expressed genes (DEGs) common to both datasets showed similarity between the 28- and 92-day transcriptome datasets (Nault *et al*., 2016). Conversely, our liberality-based analysis revealed differences, aligning more closely with observed clinical and pathological changes. Analyses comparing pathways enriched among DEGs specific to short- and long-term exposure models would help identify transcriptional transitions during disease progression (Yeh *et al*., 2026). However, such analyses offer only a partial view of the underlying transcriptomic perturbation and rely heavily on existing biological knowledge and annotation databases. Liberality, a metric that does not require *a priori* biological annotation, could provide a more objective foundation for integrating transcriptomics into chemical risk assessment.

In conclusion, the liver transcriptomes of mice exposed to TCDD showed dose-dependent increases in liberality. Overall, comparative analysis of liberality clearly distinguished the overall transcriptomic impacts of 28- and 92-day TCDD exposures. Additional case studies are needed to explore the broader relationship between liberality and adversity.

## Acknowledgements

We would like to thank Dr. Norichika Ogata for providing valuable insights into the concept of liberality and for guidance on model comparison using R. We thank ENAGO for editing a draft of this manuscript.

